# Neuropeptide Receptors Harbor More Genomic Regulatory Elements in Their Loci Than the Loci of Their Ligands

**DOI:** 10.64898/2026.05.26.727860

**Authors:** Hadi Najafi, Seung Hun Jeong, Woo Jae Kim, Sun Shim Choi

**Author notes:** **Correspondence to:** Woo Jae Kim & Sun Shim Choi. **Contact information:** HN, SHJ.

## Abstract

The neuropeptide signaling pathway is vital for the physiology and behavior of multicellular organisms. This pathway is mediated by ligand-receptor binding, wherein neuropeptides (NPs) are often released from neurons, while their receptors (NPRs) are ubiquitously expressed in both neuronal and non-neuronal cell types. To elucidate the underlying mechanisms of this expression pattern divergence, here we dissect the genomic and epigenomic architectures of these two distinct gene classes.

We first characterized the genomic architecture of NP and NPR genes to compare their total gene length, exon and intron lengths, exon counts, and number of alternatively spliced transcripts per gene. We also profiled their regulatory genomic elements including CpG islands, TATA boxes, and their overlapping antisense transcripts. These analyses were then expanded to non-human model organisms to evaluate the evolutionary conservation of the underlying mechanisms.

We found that NPRs encompass larger genomic loci and encode longer transcripts than NPs. Furthermore, the increased length of NPR transcripts was driven by longer exons rather than higher exon counts. Consistently, the number of spliced variants per gene was similar between NPs and NPRs, suggesting that alternative splicing events are a minor contributor to their distinct expression patterns. At the RNA level, NPR mRNAs possess significantly longer 3ʹUTRs compared to NPs, indicating a greater potential for post-transcriptional gene regulation. This complexity is also mirrored at the chromatin level, where NPR loci exhibit a higher density of epigenetic marks than NPs. Together, these findings highlight the multi-layered nature of gene regulation prioritizing control at the receptor level.

## Introduction

Neuropeptides (NPs) are small proteins synthesized and released by most major types of neurons that act as neurotransmitters or neurohormones to modulate diverse physiological functions—such as metabolism, emotional states, pain, locomotor activity—as well as behavioral characteristics like social interaction, feeding, and aggression (1). NPs execute their functional roles by binding to their cognate receptors—typically high-affinity transmembrane proteins such as G-protein-coupled receptors (GPCRs)—which are expressed in both neuronal and non-neuronal cell populations (2). This binding initiates an intracellular signaling cascade that propagates to mediate a wide range of cellular processes that finally affect an organism’s development, physiology, and behavior (3, 4).

The structural motifs of NP and NPR proteins, as well as their genomic architecture at the nucleotide and protein levels, are conserved across diverse taxa (5, 6). This evolutionary stability underscores their fundamental roles in metazoan life and validates the use of diverse model organisms to gain insights into their functional and regulatory mechanisms. Despite extensive research into their structures, expression patterns, and the phenotypic effects of their dysregulation, a comprehensive analysis of NP and NPR genomic features and the conserved upstream mechanisms governing them is still lacking.

Moreover, aberrations in the expression levels of NP and NPR genes are associated with various diseases including but not limited to neurodegenerative and neuroimmune diseases (7), psychiatric disorders (8), cognitive impairments (9). Furthermore, these aberrations contribute to the pathogenesis of systemic diseases beyond the nervous system, such as heart and kidney failure (10). These diverse roles of NP and NPR signaling—collectively referred to as the ‘neuropeptidergic’ signaling pathway (3, 11) reflect its fundamental importance to human health.

Mechanistic study of NP and NPR genes in human specimens faces significant limitations, primarily due to ethical concerns and the difficulty of tracing pathophysiological and behavioral traits following the over- or down-regulation of genes in live subjects. To circumvent these issues, simple model organisms—such as *Drosophila melanogaster* (*D. melanogaster*), *Caenorhabditis elegans* (*C. elegans*), *Mus musculus* (mouse), *Rattus norvegicus* (rat)—are invaluable. These organisms facilitate *in vivo* mapping of RNA and protein distribution, physiological and behavioral assays, and developmental tracing under various NP/NPR expression levels.

Research utilizing model organisms to study NP/NPR signaling has revealed that NP expression is primarily restricted to neuronal cells, whereas their cognate receptors are often widely distributed across both neuronal and non-neuronal cells (12). At the functional level, technologies such as optogenetics (13, 14) and calcium imaging (15, 16) facilitate the tracing of neural dynamics with cellular resolution and high temporal precision—capabilities typically restricted to non-human model organisms (17).

Notably, the activity and pattern of neuropeptidergic signaling pathways are strictly contingent upon the expression pattern of both NP and NPR genes. If an NP is secreted in the absence of sufficient receptor expression, or if the receptor is expressed ectopically, the whole network’s activity is disrupted (18). Conversely, if an NPR is expressed in a target tissue without the corresponding release of its NP ligand, the signaling system remains functionally silent. Thus, the efficacy of the pathway relies on the precise spatiotemporal overlap of both components. For instance, the activity of CCHamide-2—an NP produced and released by the sensory neurons within the *Drosophila* gut—is highly dependent on the expression level and distribution of its receptor. It has been demonstrated that the *Drosophila* brain expresses high levels of the CCHamide-2 receptor, whereas the gut lacks significant expression. This disparity leads to tissue-specific signaling activity due to the differential expression patterns of the receptor (19).

Despite these findings, it remains a critical objective to comprehensively determine which component of the neuropeptidergic signaling axis—the ligand (NP) or the receptor (NPR)—serves as the primary regulatory bottleneck. Clarifying these regulatory dynamics allows both basic researchers and clinicians to identify the primary ‘driver’ of the pathway. This is essential for elucidating the etiology of neuroendocrine disorders and for pinpointing high-priority therapeutic targets in human diseases driven by NP or NPR dysregulation.

In this study, we exploit the availability of the genome sequences and their multi-layered regulatory elements across four model organisms, alongside the human genome, to determine the primary locus of control for neuropeptidergic signaling regulation. By adopting a comparative approach, we aim to provide mechanistic insights into ligand-receptor signaling regulation.

## Results

### Differential genomic region configuration between neuropeptides (NPs) and neuropeptide receptors (NPRs)

While neuropeptides (NPs) are known to be produced and released by restricted cell types (primarily neurons), their receptors (NPRs) are widely expressed across an organism’s body, including neuronal and non-neuronal cells (3). Given this widespread expression pattern, we hypothesized that NPR genomic loci might be more enriched for regulatory elements than NPs, allowing them to act as substrates for regulatory processes across various cellular conditions. To test this hypothesis, we analyzed the genomic loci of these genes and found remarkable differences in their structures:

NPR loci are longer and possess remarkably larger intronic regions (Fig. 1A). NPRs also encode longer transcripts (Fig. 1B), primarily due to increased exon length rather than higher exon counts. Consistent with the similar exon counts observed between NP and NPR genes, alternative splicing patterns—as indicated by the number of transcripts per gene—showed similar levels for both groups (Fig. 1C–E). The most distinctive feature of NPRs was their intronic regions, which are significantly longer than those of NP genes. This could suggest a greater potential for NPR genes to harbor genomic regulatory elements within introns (Fig. 1F).

**Figure 1.**
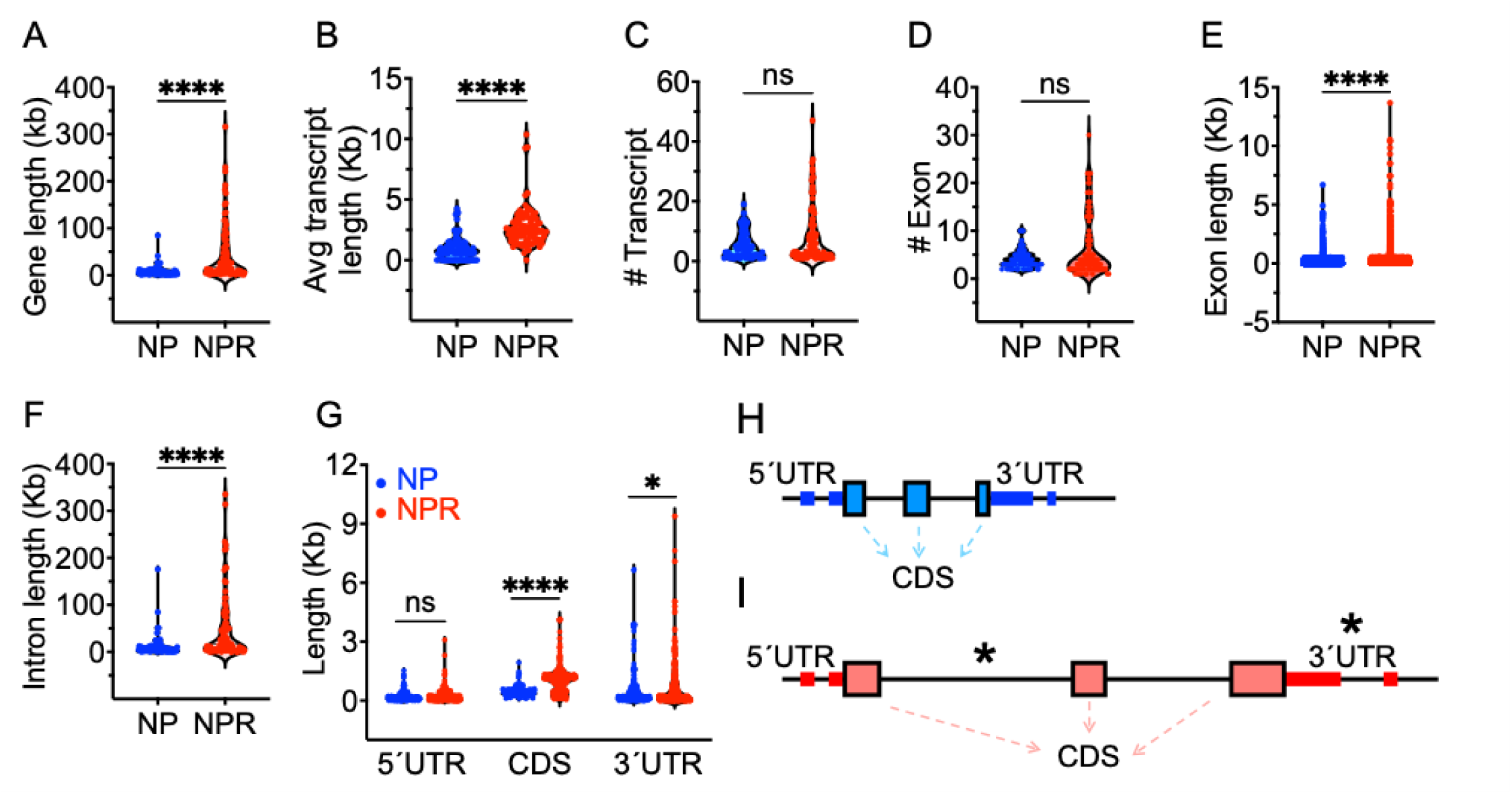
Differential genomic architecture of neuropeptide (NP) and neuropeptide receptor (NPR) genes. **A)** Increased gene length (kb, kilobase pair) in NPRs compared to NPs. **B)** NPR genes encode longer transcripts than NP genes, as indicated by the average length of all encoded transcripts per gene. **C)** The average number of transcripts encoded by each NP and NPR gene is similar, indicated by a non-significant (22) difference. **D)** NP and NPR genes exhibit similar exon counts. **E)** Average exon length is significantly greater in NPR genes than in NPs. **F)** NPR genes possess significantly larger intronic regions compared to NP loci. **G)** Analysis of mRNA features shows longer lengths for both coding sequences (CDS) and 3ʹ-untranslated regions (3ʹUTRs) in NPR mRNAs, which are critical for their proper protein function and regulatory processes, respectively. **H)** Schematic representation of the differential genomic structures of NP (blue) and NPR (red) genes. Intergenic and intronic regions are shown as solid black horizontal lines; protein-coding exons (CDS) are shown as boxes; and non-coding exons (5ʹUTRs and 3ʹUTRs) are shown as thick horizontal lines. Asterisks highlight enriched regulatory sites differentially within the introns and 3ʹUTRs of NPR genes.

When the messenger RNAs (mRNAs) of these genes were analyzed, we observed that the CDS regions of NPR mRNAs exhibited significantly greater lengths (as expected, due to their longer protein sequences), followed by significantly longer 3ʹUTRs, compared to the corresponding regions in NP mRNAs (Fig. 1G). Given the importance of the regulatory role of 3ʹUTRs (20, 21), the increased 3ʹUTR lengths in NPR mRNAs suggests a higher capacity for regulation at the post-transcriptional level—compared to their NP counterparts—such as mRNA degradation or translation inhibition by microRNAs (miRNAs).

These findings highlight a unique property of NPR genes compared to NPs: the presence of significantly longer DNA elements, such as introns, and longer RNA regulatory domains, such as 3ʹUTRs (Fig. 1H). This makes NPRs ideal substrates for regulation at the genomic and transcriptomic levels.

### Enhanced evolutionary constraint and tighter selective control over NPRs versus NPs

Detailed mechanistic analysis of NP and NPR genes necessitates the use of model organisms to trace molecular, cellular, and organismal phenotypes in systems such as mice, rats, flies, and nematodes. To evaluate the evolutionary conservation patterns of NP and NPR proteins, we utilized the DIOPT database to identify human NP and NPR orthologs across these four model organisms. Our results revealed that a higher percentage of human NPR genes possess orthologs in the evolutionarily divergent organisms (e.g., flies and nematodes), compared to NP genes (Fig. 2A–B). Consistently, dN/dS analysis between the human and mouse genomes revealed a significantly lower rate of non-synonymous amino acid changes (i.e., lower *dN/dS* ratio) in NPRs compared to NP genes (Fig. 2C). These data reflect the stronger influence of purifying selection on NPRs relative to NPs.

**Figure 2.**
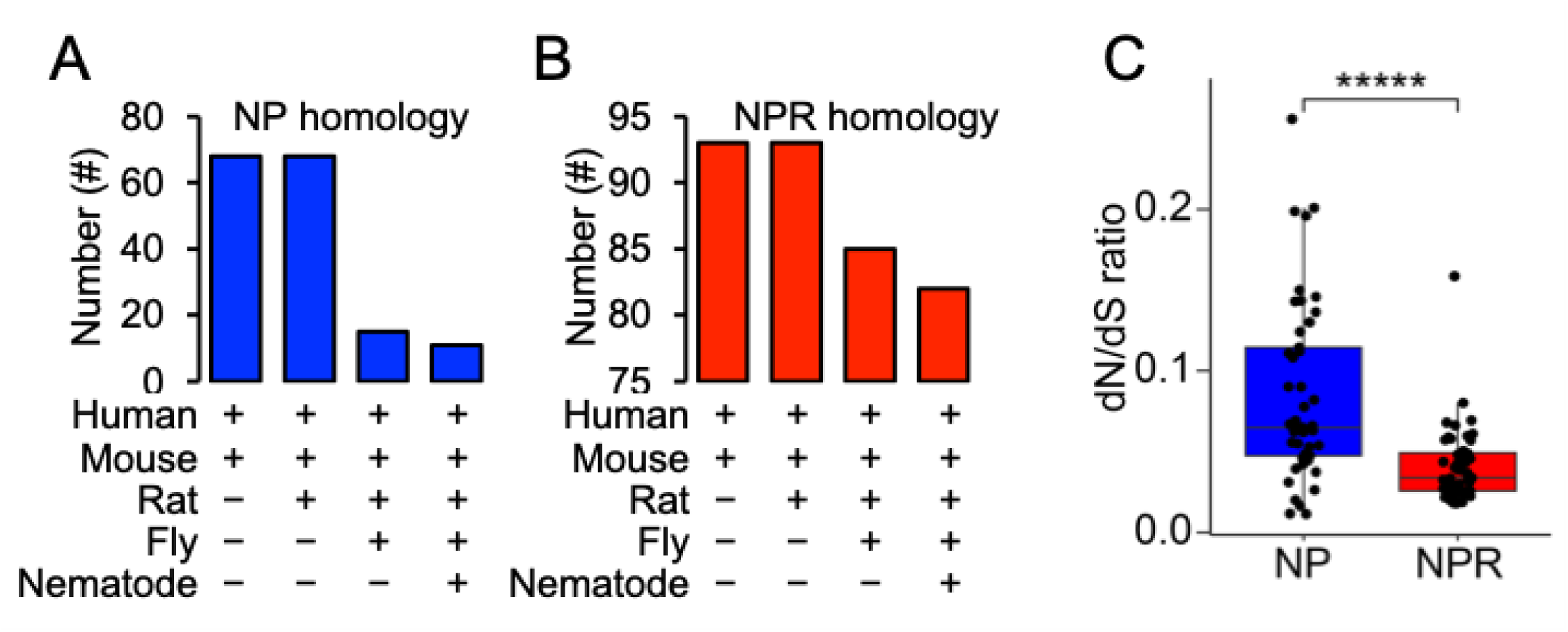
Evolutionary conservation of neuropeptide (NP) and neuropeptide receptor (NPR) genes. **A)** Number of human ‘NP’ orthologs identified across diverse species. **B)** Number of human ‘NPR’ orthologs identified across diverse species. **C)** Comparison of non-synonymous substitution rates in NP and NPR proteins between human and mouse genomes.

### Unique structural features of NPR genes are evolutionarily conserved

Given the relatively simple nervous systems and well-established neuronal connectomes of model organisms, (23), identifying the key structural features of NP and NPR genes across these species is essential.

We investigated whether the architecture of NPR genes in non-human organisms exhibits the same unique characteristics as those found in humans. We found that across all the studied model organisms—mouse, rat, fly, and nematode—NPR genes encompass longer genomic regions (Fig. 3A–B). Similarly, the average lengths of NPR transcripts were consistently greater than those of NP transcripts, suggesting that the structural distinctions between these two gene families are conserved from humans to insects and nematodes (Fig. 3C–D). Like their human counterparts, the NP and NPR genes of these model organisms did not show a significant difference in their ‘number of transcripts per gene’, suggesting comparable levels of alternative splicing event for both gene classes (Fig. 3E–F).

**Figure 3.**
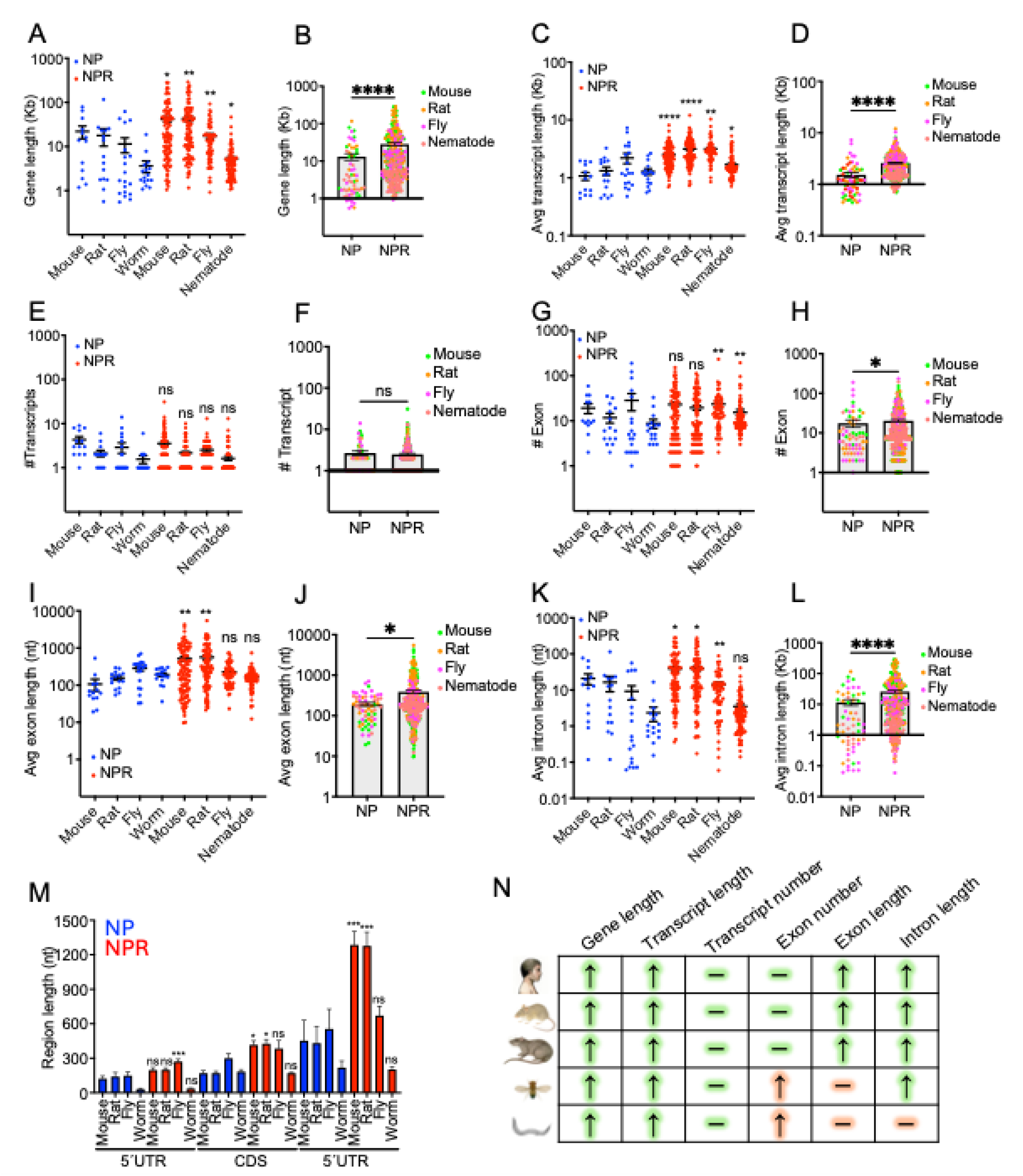
Genomic and transcriptomic features of neuropeptide (NP) and neuropeptide receptor (NPR) genes across model organisms. **A)** Comparison of gene lengths, shown on a log-transformed scale (kb, kilobase pair) for each organism. **B)** Average gene lengths of NP and NPR genes, representing the cumulative mean differences across all studied organisms. **C)** Comparison of transcript lengths, shown on a log-transformed scale (kb) for each organism. **D)** Average transcript lengths of NP and NPR genes, representing the cumulative mean differences across all studied organisms. **E)** Comparison of the ‘number of transcripts per gene’ for NPs and NPRs of each organism. **F)** Cumulative representation of number of transcripts per gene across all studied organisms. **G)** Comparison of the ‘number of exons per gene’ for NPs and NPRs of each organism. **H)** Cumulative representation of the ‘number of exons per gene’ across all studied organisms. **I)** Comparison of exon lengths, shown on a log-transformed scale (nt, nucleotides) for each organism. **J)** Average exon lengths of NP and NPR genes, representing the cumulative mean differences across all studied organisms. **K)** Comparison of intron lengths, shown on a log-transformed scale (22) for each organism. **L)** Average intron lengths of NP and NPR genes, representing the cumulative mean differences across all studied organisms. **M)** Average lengths of mRNA features (5ʹUTR, CDS, and 3ʹUTR) across all studied organisms. Asterisks represent unpaired t-test comparisons of the NPR group (red bars) with its NP counterpart (blue bars) within each organism. **N)** Overall pattern of gene structural features across all model organisms included in this study. Green highlights represent fully evolutionarily conserved features, while red highlights indicate exceptions for the features observed in the fly and nematode. Human: *Homo sapiens*, Mouse: *Mus musculus*, Rat: *Rattus norvegicus*, Fly: *Drosophila melanogaster*, and Nematode: *Caenorhabditis elegans*.

Similar to the longer exon lengths seen in human NPRs, NPR genes in mouse and rat exhibited a longer average exon length, accounting for their longer transcripts. Also, mouse and rat showed no significant differences between the exon counts of NP and NPR genes. However, in fly and nematode, NPR genes exhibited higher exon counts despite having similar average exon lengths (Fig. 3G–J), indicating that longer transcripts in these species are driven by exon number rather than exon size.

Except for nematode, all organisms exhibited a significant increase in the length of the intronic regions of NPR genes in comparison with those of NPs (Fig. 3K–L). Despite this exception, a cumulative comparison of all studied genomic features showed a similar trend across humans and model organisms, reflecting a relatively conserved pattern of NP and NPR gene structures across species (Fig. 3B–L, bar graphs).

Both the CDS and 3ʹUTR in rodents (mouse and rat) exhibited significantly longer lengths in NPRs, while 5ʹUTR lengths remained similar— a pattern consistent with humans. In contrast, no such differences were observed in the nematode or the fly, with the sole exception of the fly’s NPR mRNAs, which exhibited longer 5ʹUTRs compared to NP mRNAs (Fig. 3M).

Altogether, these analyses demonstrate that NP and NPR gene structures are largely conserved across species (Fig. 3N, green highlights). The primary deviations from this pattern are the exon counts and lengths in *D. melanogaster* (fly) and *C. elegans* (nematode), alongside unique intron length variations in nematode (Fig. 3N, red highlights).

### NPR genomic loci are subject to more epigenetic modifications than NP loci

We showed that NPR genes occupy larger genomic loci and exhibit a distinct architecture of longer exons and introns (Figs. 1 and 3). Given that these features are metabolically costly to transcribe and process as heterogeneous nuclear RNA (hnRNA) products, we hypothesized that NPR expression must rely on robust regulation at the ‘genomic’ level. We propose that they utilize a more diverse array of epigenetic marks than NP genes to manage this regulatory demand efficiently.

To test this hypothesis, we investigated the levels of major well-characterized epigenetic marks, including signatures of active promoters (H3K4me3), active enhancers (H3K27ac and H3K4me1), active transcription (H3K36me3), and gene silencing (H3K27me3 and H3K9me3). Utilizing ChIP-sequencing (ChIP-seq) datasets from ENCODE, we examined these epigenetic signatures across three human cell lines—human embryonic stem cells (hESCs), neural stem cells (NSCs), and fully differentiated neurons—representing different developmental stages.

Results highlighted a distinct epigenetic signature across the promoter and gene body regions of NPRs compared to NPs. Specifically, NPR genes harbor more regulatory elements for promoter activation (elevated H3K4me3 signal) and exhibit more bivalent modifications on their gene bodies, corresponding to alternating transcriptional elongation (ON) and chromatin suppression (OFF) states (Fig. 4A, *P* < 0.05). These data suggest that NPR genes possess a higher potential than their NP counterparts to serve as substrates for complex regulatory processes by epigenetic modifiers (Fig. 4B).

**Figure 4.**
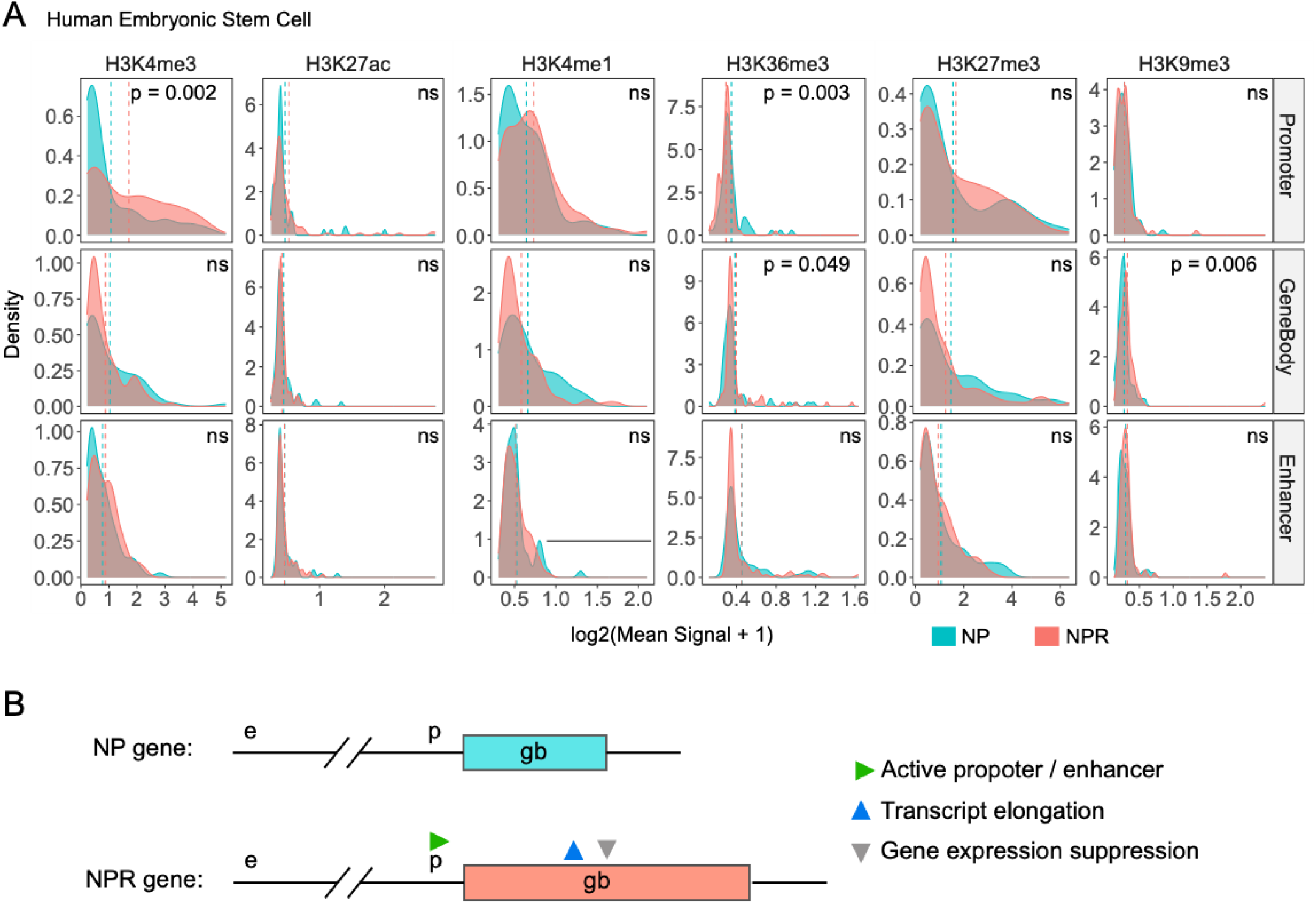
Distinct profile of epigenetic marks on neuropeptide (NP) and neuropeptide receptor (NPR) genomic loci. **A)** ChIP-seq analysis of major functional epigenetic modifications across three genomic regions (promoter, gene body, and enhancer) in human embryonic stem cells (hESCs): The promoter regions of NPR genes (red) exhibited significantly higher epigenetic signals compared to those of NP genes (24). Within the gene body, signals associated with transcriptional elongation and gene silencing were also higher in NPRs than in NPs. A p-value < 0.05 was considered statistically significant based on the average signal frequencies across samples; data are presented on a log-transformed scale, log2(Mean+1). **B)** Schematic representation of enriched epigenetic modification sites (triangles) in NPR genes relative to NP genes, facilitating adaptable and fine-tuned expression levels. The promoter region (’p’) of NPR genes harbors more activation sites (increased H3K4me3, green triangle), while the gene body (’gb’) exhibits bivalent regulatory potential, containing both transcriptional activators (H3K36me3, blue triangle) and silencers (H3K9me3, grey triangle). e = enhancer, p = promoter, and gb = gene body.

To assess whether these epigenetic patterns are maintained across human neuronal lineages with varying differentiation states, we extended our analysis to neural stem cells (NSCs) and fully differentiated neurons using ENCODE ChIP-seq datasets (Supplemental file S2A).

Consistently, NPR genes exhibited higher signal distribution for the active promoter mark (H3K4me3), as well as increased transcriptional elongation (H3K36me3) and repressive (H3K9me3) marks, compared to NP genes across both promoter and gene body regions (Fig. 5A and 5B).

**Figure 5.**
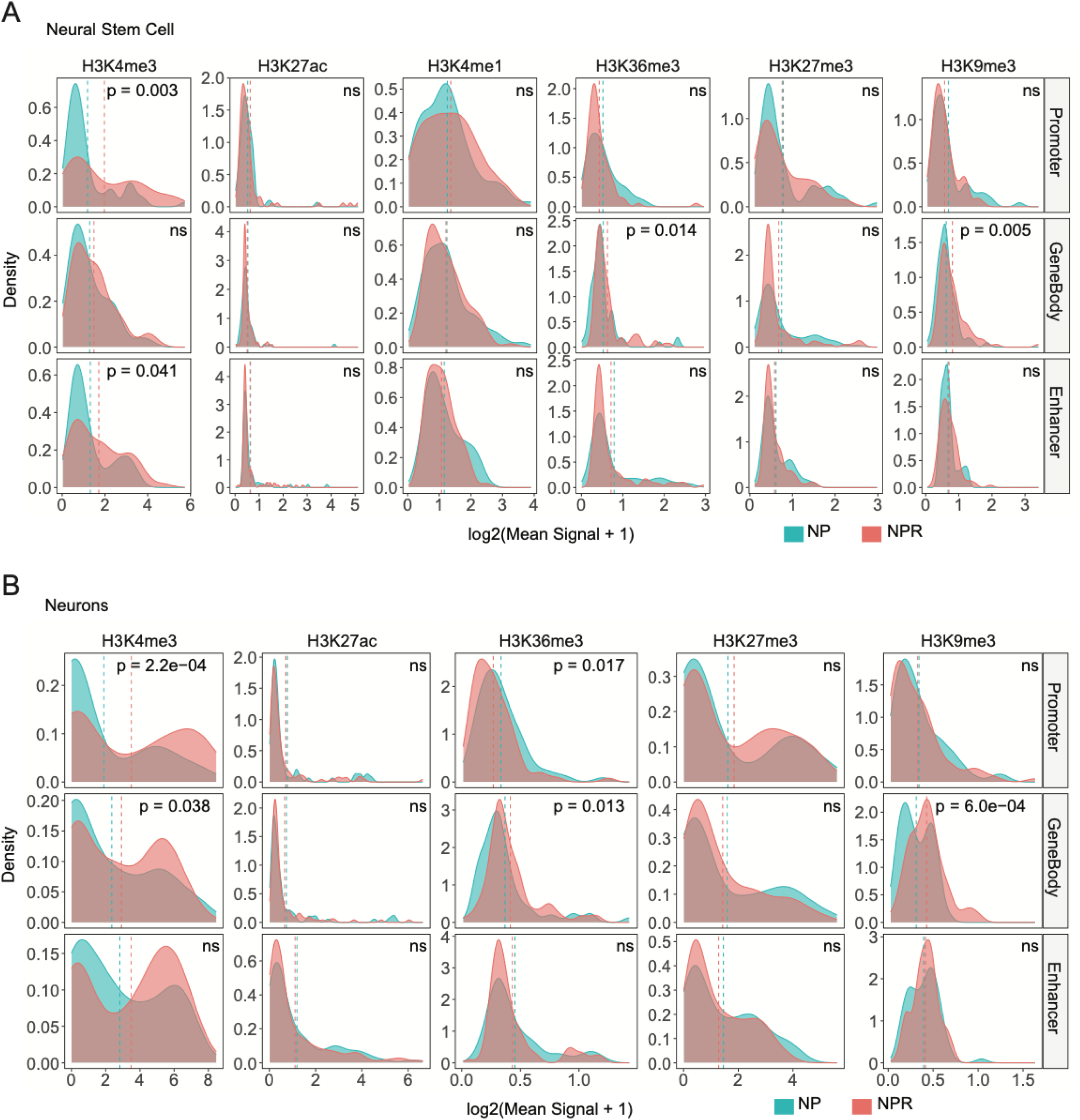
Sustained differential epigenetic profiles of NPR genes during neuronal cell differentiation. **A)** In neural stem cells (NSCs), NPR genes (red) exhibit higher epigenetic signal distributions compared to NP genes (blue) across promoter and gene body regions, with significant differences observed for selected histone marks. **B)** In fully differentiated cells (neurons), these elevated epigenetic signals in NPR genes are maintained, indicating sustained regulatory differences during neuronal differentiation. Differences between NP and NPR groups were assessed using the Wilcoxon rank-sum test (unpaired), following assessment of non-normal data distribution. Data are presented as log-transformed signal values (log2(Mean signal + 1)). A P-value < 0.05 was considered statistically significant.

These findings indicate that NPR genes possess more dynamic regulatory features, acting as substrates for both transcriptional activation and repression within the same genomic context. This regulatory flexibility is likely associated with the fine-tuned expression requirements of NPR genes during neuronal development and function.

### NPR genes are often localized within domains of high chromatin dynamism

To survey the state of chromatin dynamics across human NP and NPR genes, we utilized public ChIP-seq datasets from ENCODE and analyzed these data for epigenetic modifications representing four distinct chromatin states: active, silenced, bivalent, and inactive (Figure 6).

**Figure 6.**
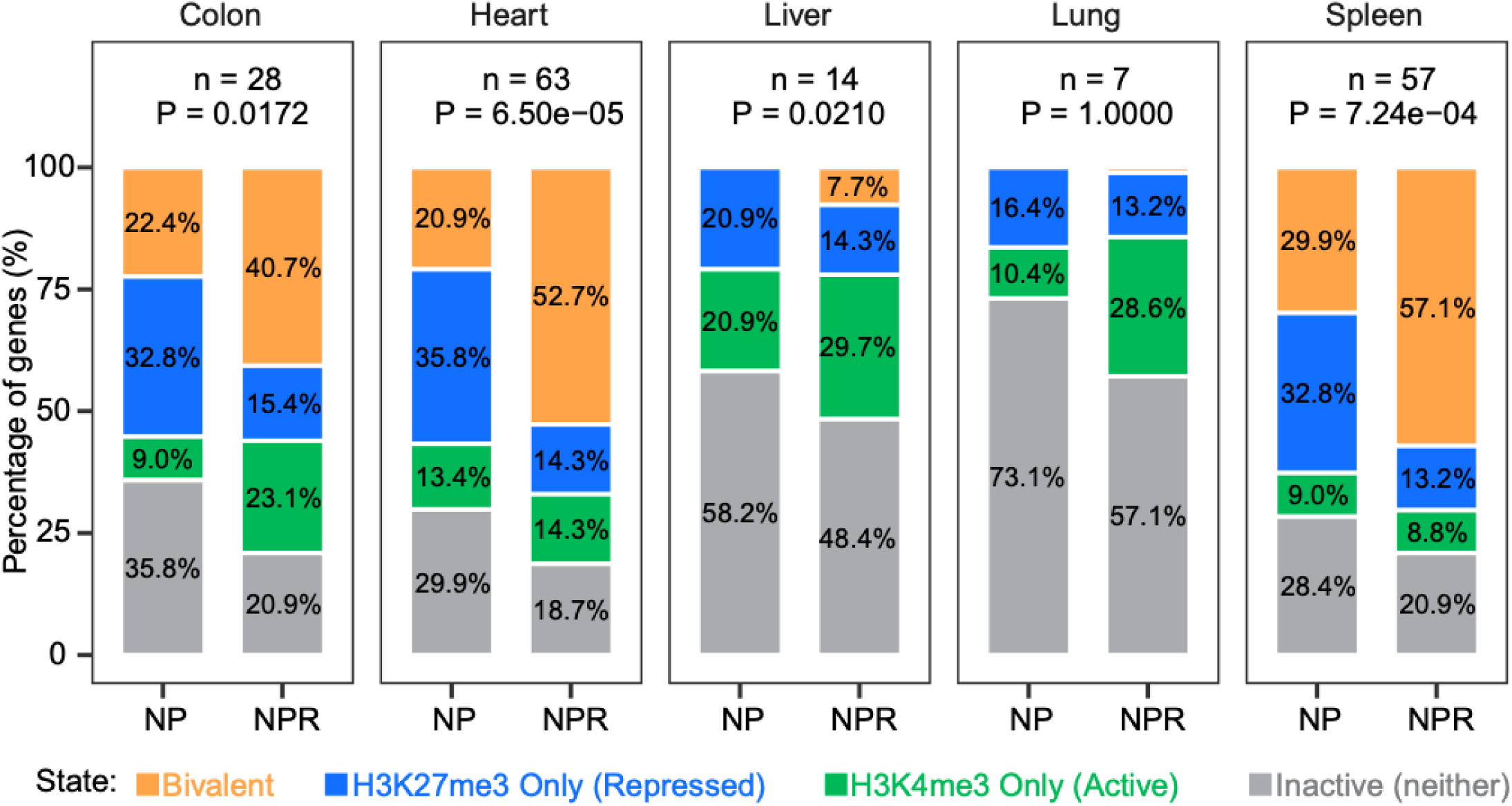
Distinct chromatin dynamics for neuropeptide (NP) and neuropeptide receptor (NPR) genes. Across nearly all the examined human tissues, a higher proportion of NPR genes harbor genomic sites characterized by bivalent chromatin (orange) and activating epigenetic modifications (24). Conversely, NPR genes consistently exhibit fewer repressive modifications (blue) or inactive chromatin states (grey) compared to NP genes. ‘n’ represents the number of ChIP-seq dataset (human specimen) for each tissue. The P-value in each graph represents the statistical significance of the distributions of epigenetic states between the two groups of NP and NPR genes. The non-significant difference in lung tissue (P=1.0000) is likely due to a smaller sample size (n=7). Despite this limitation, a similar distribution of chromatin states is observed for the lung.

Results showed a significant enhancement of the bivalent chromatin state in NPR genomic regions compared to NPs (Fig. 6, orange). These regions encompass both activating (H3K4me3) and repressive (H3K27me3) histone modifications. These data suggest that NPR genes possess greater potential than NPs to serve as substrates for epigenetic regulation, providing increased adaptability to upstream regulators.

Consistent with the broad expression patterns of NPRs—reminiscent of housekeeping genes—a smaller percentage of NPR genes harbor solely H3K27me3 repressive sites (Fig. 6, blue). This suggests fewer epigenetic constraints on NPR genes, a finding further supported by the enrichment of H3K4me3 activator modifications in NPRs relative to NPs (Fig. 6, green). Conversely, a higher proportion of NP genes lacked these epigenetic signatures entirely (Fig. 6, grey), suggesting they have reduced potential for dynamic epigenetic modification and regulation in comparison with their receptors (NPRs).

### Supportive evidence of enriched regulatory elements at NPR loci compared to NP loci

To examine whether the genomic loci of NP and NPR genes differ in harboring key transcriptional regulators—such as CpG islands, TATA boxes, and overlapping antisense transcripts—we analyzed their genomic regions for these elements in the UCSC Genome Browser. As a result, we found that human NPR genes are significantly more likely to be in proximity to, or overlapping with, other transcripts in a reverse orientation compared to their NP counterparts. These overlapping ‘antisense transcripts’ are among the most well-characterized regulators of gene expression (22, 25). Nevertheless, NPR genes in the other studied organisms did not show such a difference (Fig. 7A–E), suggesting that this genomic feature of NPR genes is human-specific.

**Figure 7.**
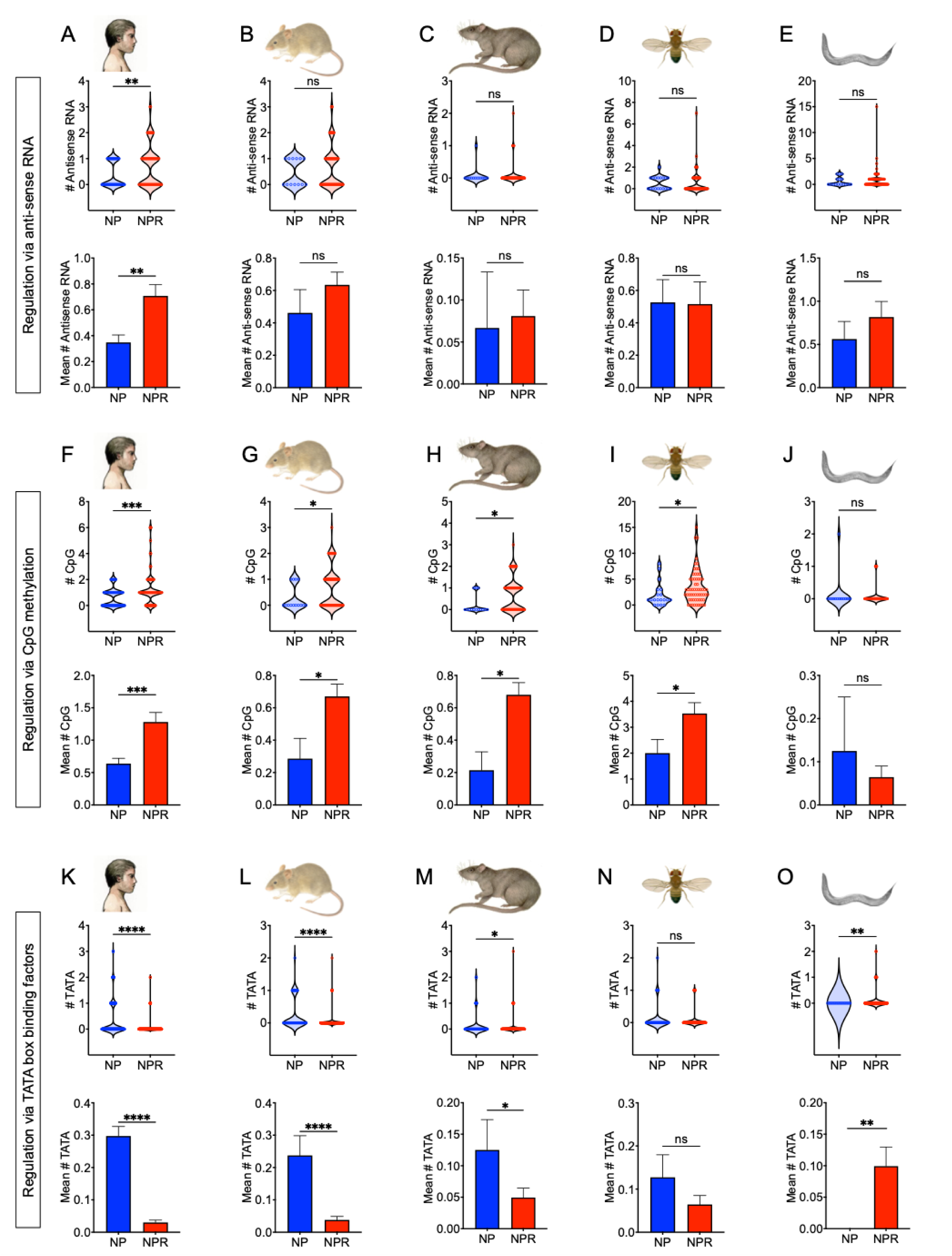
Differential distribution of key regulatory elements surrounding NP and NPR genomic loci. **A–E)** Number of antisense transcripts (RNA) overlapping with the genomic loci of NP (blue) and NPR (red) genes. Violin-dot plots illustrate the distribution of this quantification, and bar graphs represent mean value comparisons and statistical significance. **F–J)** Number of CpG islands upstream of the genomic loci of NP (blue) and NPR (red) genes. Violin-dot plots illustrate the distribution of this genomic feature, and bar graphs represent mean value comparisons and statistical significance. **K–O)** Number of TATA boxes upstream of the genomic loci of NP (blue) and NPR (red) genes. Violin-dot plots illustrate the distribution of this genomic feature, and bar graphs represent mean value comparisons and statistical significance. P<0.05 was considered statistically significant; ‘ns’ denotes non-significant differences; ∗P<0.05, ∗∗P<0.01, ∗∗∗P<0.001, and ∗∗∗∗P<0.0001.

Analysis of CpG island presence within the loci of the five organisms in this study showed that NPR genes harbor a greater number of these genomic elements compared to NP counterparts. The exception to this trend was the nematode *C. elegans*, which showed no significant differences in CpG counts between its NP and NPR genes (Fig. 7F–J). CpG islands are primary sites for DNA methylation and often serve as indicators of ubiquitously expressed genes (26, 27).

Conversely, TATA boxes—which act as control checkpoints for tissue- or cell-type-specific gene expression (28, 29)—exhibited a reverse pattern, in which ‘NP’ genes contained ‘more’ TATA elements upstream of their genomic loci.

Altogether, these findings are consistent with the broader expression pattern of NPR genes compared to NP counterparts, and the presence of more CpG islands and fewer TATA elements likely provides the molecular basis for this breadth. Unlike the antisense transcript findings, the distribution patterns of CpG islands and TATA boxes around NPR loci were not human-specific, as they were also shared by mouse and rat (Fig. 7K–O).

## Discussion

Neuropeptides (NPs) are signaling molecules secreted primarily from neurons and act as neurotransmitters or peptide hormones to affect a wide range of physiological and behavioral characteristics. They exert their functions by binding to their receptors, termed ‘neuropeptide receptors’ (NPRs), which, compared to NPs, have a broader expression pattern across the body, covering both neuronal and non-neuronal cells (19, 30). The cooperative actions of both NP and NPR proteins within their signaling processes, often referred to as the ‘neuropeptidergic signaling’ (24, 31), are essential for controlling the physiology and behavior of an organism under healthy, stressed, or diseased conditions. Therefore, understanding the molecular mechanisms governing the dynamic and adaptive activity of the NP/NPR system is necessary in the basic biology and medicine, yet this understanding has so far remained incomplete.

In this study, we dissected the genomic anatomy of the pathway’s core players: the NP and NPR genes. We found that NPR genomic regions harbor more regulatory elements, making them the ideal regulatory substrate for upstream regulatory processes. NPR genes, in comparison with their ligand counterparts (NPs), possess remarkably longer intronic regions and 3ʹ-untranslated regions (3ʹUTRs)—both known primarily as regulatory sites in the genome and transcriptome, respectively.

It was expected that NPR genes would have longer mRNA transcripts due to the necessity of longer coding sequence (CDS) regions to encode the proteins required for integration into the cell membrane. However, the observed longer 3ʹUTRs for these genes cannot be ignored. Studies have shown the significance of 3ʹUTRs in gene expression and function at the messenger RNA (mRNA) level; the 3ʹUTR serves as a docking site for RNA-interacting proteins (RBPs) that drive diverse processes, such as mRNA localization, stability regulation, and scaffolding for protein-protein interactions (32, 33). Furthermore, besides providing regulatory sites for RBPs (34, 35), 3ʹUTRs are also the primary site for the regulatory microRNAs (miRNAs) (20).

This evidence highlights the greater capacity of NPR genes, compared to their ligands, for regulation at both the genomic and transcriptomic levels. This concept aligns with established principles of gene regulation, whereby pathways with the lowest metabolic cost are evolutionarily selected (36–39). In fact, NPR genes encompass a significantly larger genomic loci (Fig. 1A and 3A–B); if their regulation were to occur primarily post-transcriptionally, the cell would incur a high metabolic cost to fine-tune the levels of their transcripts or proteins via RNase/protease cleavage or modifications (e.g., methylation, phosphorylation, acetylation, and glycosylation). Therefore, to avoid these complex and energy-consuming processes, NPR genes are evolutionarily constrained to be primarily controlled at the transcriptional level. Furthermore, while larger genomic loci incur a greater metabolic cost for transcription to the cell, their non-coding transcribed regions (e.g., introns and 3ʹUTRs) can also serve as the sites for additional functional RNAs. These include sense-overlapping genes, such as miRNAs (40), which may have functional roles either related or unrelated to their host genes, NPRs (41, 42). For example, the human NPR gene *GPR39* (ID: *ENSG00000183840*) encompasses a genomic locus of ∼229.8 kilobase pair (kb) but possesses two exons separated by a very long intronic region wherein the miRNA-9986 is in the same strand orientation (hg38; chr2:132416805-132646582; positive strand). Similarly, the *CALCR* gene (ID: *ENSG00000004948*), with a genomic locus of 150.2 kb, harbors two miRNA genes—miRNA-653 and miRNA-489—both located in its 13^th^ intron in the same strand orientation (hg38; chr7:93424486-93574724; minus strand). Due to this shared strand orientation, the miRNA loci are co-transcribed with their host genes (NPRs), hence, they may become functional once the NPR genes are transcriptionally active. These observations may explain why NPR genes have larger genomic loci, which accommodate more regulatory elements and encode additional functional products like miRNAs (42, 43).

Across the model organisms examined in this study, the genomic architectures of NP and NPR genes remain distinctly conserved (Figs. 1 and 3). The significantly longer intronic regions observed in NPR genes raised the question of whether they uniquely harbor additional gene regulatory elements compared to NPs.

To test this, we analyzed epigenetic profiles across NP and NPR genes, focusing on marks representing active promoters (H3K4me3), active enhancers (H3K27ac and H3K9me1), active transcription (H3K36me3), and silenced chromatin (H3K27me3 and H3K9me3) (44, 45). ChIP-seq analysis revealed significantly enriched signals for these marks within NPR genomic loci (Fig. 4A). This is illustrated in a schematic (Fig. 4B) showing elevated promoter activity, increased transcriptional elongation, and higher levels of silenced chromatin in NPR genes. This pattern was consistently observed across three human cell lines representing distinct developmental stages: totipotent (hESC H1), multipotent (neural stem cell, NSC), and fully differentiated (neuron) cells (Figs. 4 and 5).

Epigenetic regulation—mediated by diverse chromatin remodelers and modifiers such as methyltransferases, demethylases, Polycomb Repressive Complexes (PRC1/2), histone deacetylases (HDACs), and histone acetyltransferases (HATs)—is critical in the control of gene expression (46). These factors modulate transcription primarily by altering the topological structure and accessibility of chromatin (47), thereby providing an energetically efficient mechanism for gene regulation (i.e., preferentially at the transcriptional level). Consistent with these findings and given the enrichment of epigenetic marks at NPRs compared to their NP counterparts (Figs. 4 and 5), we observed that NPR genes possess higher levels of bivalent chromatin. Furthermore, they exhibit a distinct profile characterized by a higher prevalence of active chromatin states and a lower prevalence of repressed states across their loci (Fig. 6). Epigenetic modifications can be connected to the environmental conditions of an organism (48). Therefore, the heightened susceptibility of NPR loci to these modifications suggests that NPRs, rather than NPs, serve as the primary substrates for organismal adaptation. Together, these findings provide a rationale for the genomic architecture of NPR genes while linking these features to their essential functional roles (49).

Most human NP and NPR genes have homologs in other organisms, making these genes excellent candidates for study in popular model organisms. In this study, we performed a cross-species validation of NP/NPR genomic structures; however, significant work remains to identify the complete set of these genes in non-human organisms. Despite the conserved genomic structure of these genes across species, minute differences may be responsible for species-specific regulation of NPR genes. For instance, while NPR transcripts are consistently longer than NP transcripts across all species, as demonstrated in this study, the exon/intron architecture and the ratio of untranslated region (UTR) to the coding sequence (CDS) vary by organism. Specifically, the longer transcripts found in mammals are primarily due to increased exon length rather than higher exon counts. In contrast, in *D. melanogaster* and *C. elegans*, NPR genes exhibit higher exon counts than NP genes, while their average exon lengths remain similar (Fig. 3). Moreover, the 3ʹUTRs of NPR mRNAs in these two organisms did not trend longer than NP 3ʹUTRs (Fig. 3M). These observations could reflect: 1) Incomplete knowledge regarding the full nucleotide sequences of NP and NPR transcripts. 2) Possible distinct post-transcriptional regulatory paradigms in flies and nematodes compared to mammals.

In other words, such slight differences (e.g., exon lengths/counts, and 3ʹUTR lengths) stem in part from the inherent differences in genomic architecture across species (the biological component) and in part from a lack of sufficient knowledge (technical issues) about their transcriptome. For example, many NP and NPR genes in non-human organisms are proposed based on protein sequence homology and reverse-mapping of identified peptides to the genome; hence, their mRNAs are often reported as merely a CDS—without the precise termini of the transcripts. For example, the *W05H5.1* gene in *C. elegans* (WormBase: *WBGene00012283*, a G-protein-coupled receptors family 1 profile domain-containing protein) is reported without specifying its exact mRNA termini (CDS = 963 nucleotides, 5ʹ/3ʹUTRs = 0 nucleotides). Similarly, the *D. melanogaster CG3216* gene (FlyBase: *FBgn0034568*, a homologue for human *NPR1* gene) is reported without UTR length specification (CDS = 3291 nucleotides, 5ʹ/3ʹUTRs = 0 nucleotides).

Such technical issues are more common in non-human organisms, as most of the research has focused on human specimens (50–53), hence, the human-derived transcripts are more likely to be covered by many studies and released as their full-length sequences, possessing full sequences of all mRNA portions (i.e., complete 5ʹUTRs, CDS, and 3ʹUTRs). Experimental techniques such as ‘rapid amplification of cDNA ends’ (RACE) (54, 55), along with more advanced methods like long-read RNA sequencing (56, 57) and *in silico* gene modeling (58, 59), are necessary to identify or verify the full-length versions of these mRNA regions in the poorly characterized genes of non-human organisms.

Similarly, in addition to the noncoding portions of NP/NPR mRNAs (i.e., UTRs), the full profile and sequences of ‘intergenic’ long noncoding RNAs in the non-human genomes are also incomplete. Consistent with this fact, we found that NPR genes are more frequently located at genomic loci neighboring antisense overlapping transcripts; however, this trend was not observed in non-human organisms, such as mouse, rat, fly, and nematode (Fig. 7A–E). This discrepancy might originate from the unique composition of the human genome—for example, the specificity of noncoding RNA prevalence in more advanced organisms like primates versus simpler organisms like flies and nematodes (60). Long noncoding RNAs (lncRNAs)—unlike coding ones (i.e., mRNAs)—are considered the ‘dark matter’ of evolution (61, 62). They are significantly more prevalent in higher primates compared to simpler organisms such as rodents, insects and nematodes (63), and may lack significant evolutionary conservation at the nucleotide sequence level (64).

Since these transcripts do not encode proteins and are not evolutionarily conserved, they cannot be easily discovered via computational predictions or genome modeling. Therefore, more effort in the discovery and functional verification of these transcripts is required for non-human model organisms (65, 66). However, in the human genome, there is a wealth of studies discovering these transcripts, sufficient to conclude that NPR genomic loci tend to be in the proximity of antisense transcripts. This makes NPRs substrates of this type of gene expression regulation at both the transcriptional (67) and post-transcriptional levels (67, 68): Antisense transcripts can match (via Watson-Crick base pairing) intracellularly with their overlapping transcripts, bringing the formed double-stranded RNA (dsRNA) to another level of gene expression regulation (68, 69). Additionally, antisense RNAs can recruit epigenetic and transcription factors to the loci of their neighboring genes, playing roles in transcriptional induction or suppression (67).

Via analysis of genomic regions surrounding NP and NPR genes, we found a significantly enriched presence of antisense noncoding RNAs for NPR genes versus NPs, which was specific to humans (Fig. 7A–E). Investigating the putative link between these coding-noncoding pairs (70, 71) in these loci is of great importance. Such an examination will add another layer of regulation to the neuropeptidergic signaling pathway and establish lncRNAs as a new class of master regulators for this system.

NPRs also exhibit a differential genomic architecture in terms of harboring different frequencies of two classes of transcription factor binding sites (TFBSs): CpG islands and TATA boxes. NPR genes possess significantly more CpG islands (an indication of ubiquitously expressed genes) while being depleted of TATA boxes (an indication of cell- or tissue-specific genes) in comparison with NP genomic loci. These findings are also in line with the expression distribution patterns reported in previous studies.

As an example of an NP-NPR pair, we examined the loci of *NPY* gene and its primary receptor, the *NPY1R* gene, to illustrate the structural differences between NPs and NPRs. *NPY* (one of the most studied NPs in human) exhibits a narrower expression pattern in the human body in comparison with the broader expression pattern of its receptor, *NPY1R*. Furthermore, more transcription factors (TFs) are found to bind to the *NPY1R* locus than to that of *NPY* gene. Functional classification of the TFs differentially bound to the *NPY* and *NPY1R* genes revealed that *NPY1R*-specific TFs are often classified as ‘activators’, ‘structural’ or ‘architectural’ (e.g., scaffolds and chromatin-bound factors), whereas *NPY*-specific TFs are often dual-functioning (ON or OFF mode) (Supplemental file S4A). These findings suggest crucial structural differences between NPRs and NPs. Given these differences in genomic architecture—specifically the ‘frequency’ and ‘type’ of their regulatory elements—distinct expression patterns for NP-NPR gene pairs are expected; indeed, this was consistently observed for *NPY* and *NPY1R* genes (Supplemental file S4B), showing a lack of expression correlation between *NPY* and *NPY1R* transcription levels in most human tissues. Gene set enrichment analysis (GSEA) (72) on the identified NP- and NPR-specific TFs highlighted the enrichment of the human *chromosome 20*—specifically the *chr20q13*, *chr20q11*, and *chr20p11*—as the most enriched chromosomal locations harboring the NPR-specific TFs including *CEBPB*, *CTCFL*, *TFAP2C*, *E2F1*, and *FOXA2*. However, no chromosomal enrichment was observed for the NP-specific TFs (Supplemental file S5A–B). This data emphasizes the great potential of NPR loci as a primary site of ‘pioneer’ TFs (73–75) to initiate a molecular cascade which begins with transcription from NPR genes. Further experimental studies such as ‘genome-wide association studies’ (GWAS) and ‘expression quantitative trait loci’ (eQTL), are required for finding the putative functional link between these loci on the human *chromosome 20* and neuropeptidergic signaling.

## Methods and materials

### Input data

Most recent comprehensive lists of human neuropeptides (NPs) and neuropeptide receptors (NPRs) were obtained from the IUPHAR/BPS database (76, 77). Gene ID conversion—from official gene symbols to ‘full gene names’ and Ensembl (78) gene IDs—was performed through the one-by-one searching of the NP/NPR genes in NCBI ‘Entrez gene’ database (https://www.ncbi.nlm.nih.gov/gene/) (Supplemental file S1A).

Homology search of these human gene lists was performed for identification and listing complete set of NP and NPR orthologues in other organisms commonly used as model organisms: mouse (*Mus musculus*), rat (*Rattus norvegicus*), fly (*Drosophila melanogaster*), and nematode (*Caenorhabditis elegans*). To this end, the DIOP database (https://www.flyrnai.org/cgi-bin/DRSC_orthologs.pl) was used and human Ensembl gene IDs were used as query in this database. DIOP uses the amino acid sequence of the queries like NP or NPR gene symbols or IDs to provide a comprehensive list of homologues and it covers finding homologues in a diverse set of organisms (79). All identified orthologues of human NP and NPR genes listed by DIOPT were included in this study for downstream analyses. DIOPT provides both official gene symbols and unique gene IDs of its results. These gene IDs for human, mouse, and rat are unique Ensembl gene IDs. For example, *ENSG00000254647*, *ENSMUSG00000000215*, *ENSRNOG00000020405*, are Insulin or Insulin-like peptides in human, mouse, and rat, respectively. However, gene IDs for the NP/NPR homologues in fly and nematode in DIOPT results were specific to their assigned databases—FlyBase and WormBase, respectively. For example, the gene IDs *FBgn0044050* and *WBGene00002084* are Insulin homologues in Fly and worm, respectively. The resultant converted unique gene IDs were used as query to search in UCSC (80) for analysis of the genomic features of NP/NPR genes in human and these model organisms (Supplemental file S1B). Number of overlapping antisense RNAs, and the number of CpG islands in the genomic loci of the genes were evaluated and counted from UCSC genome browser and presented for each NP and NPR gene group side-by-side in bar graphs with dots. Each dot represents the status of one NP or NPR gene for the indicated features.

For accurate measuring of gene length, number of exons/introns, number of transcripts, average transcript length, average exon and intron lengths, as well as messenger RNA (mRNA) features including 5ʹ-untranslated region (5ʹUTR), coding sequence (CDS) and 3ʹ-untranslated region (3ʹUTR) per each gene, latest update of ‘Gene Transfer Format’ (GTF) file of each organism was downloaded from Ensembl (Release 115) and analyzed. These GTF files corresponded to the last updated genome versions of these organisms including *GRCh38.p14* for human, *GRCm39* for mouse, *GRCr8* for rat, *BDGP6.54* for fly, and *WBcel235* for nematode.

### Structural analysis of genes and transcripts

For side-by-side comparison of NP and NPR gens based on their specific gene features as well as their embedded regulatory elements, the most recent release of the genome sequences for human, mouse, rat, fly and nematode together with their corresponding GTF files were used and calculations were carried out as following:

The ‘gene length’ parameter was defined as the interval between the first nucleotide of the gene and the last nucleotide of its final exon, encompassing both exonic and intronic sequences. Gene and transcript structural features were extracted from GTF annotation files using custom Python scripts. Briefly, Ensembl gene IDs corresponding to NP and NPR genes were first mapped to their official gene symbols. The GTF file was then parsed to retain entries annotated as ‘transcript’ or ‘exon’ corresponding to NP/NPR genes. For each retained feature, genomic coordinates were extracted, and the feature length was calculated as (end – start + 1). The number of transcripts per gene was determined by counting GTF entries annotated as “transcript” for each gene.

Since each gene potentially encodes at least one transcript, the ‘average transcript length’ was calculated for each NP and NPR gene. To this end, GTF files were used, and the average transcript length for each gene was calculated by dividing the sum of all its transcript lengths by the total number of transcripts encoded by that gene. A similar procedure was applied to determine the exon and intron counts, as well as their average values for length—defined as the ‘average exon length’ and ‘average intron length’ per gene.

To examine the structural features of NP and NPR mRNAs, counts and lengths of their coding (CDS) and noncoding regions (5ʹ and 3ʹ UTRs) were determined and presented using the downloaded GTF files, opened in Microsoft Excel and counted as similar approach for other gene features such as exons and introns.

### Cross-species evolutionary conservation analysis

To investigate the evolutionary conservation of human NP and NPR genes, the ratio of non-synonymous to synonymous substitutions (dN/dS) was calculated using human and mouse orthologous gene pairs. Orthologs were identified using a reciprocal best-hit (RBH) strategy based on BLAST searches, where protein-coding sequences from human and mouse genome annotation files (GRCh38.p14) were queried against each other using BLASTX.

For each orthologous gene pair, aligned coding sequences were used to estimate the number of non-synonymous (dN) and synonymous (24) substitutions. The dN/dS ratio was calculated using the KaKs Calculator. In this context, dN represents nucleotide substitutions that alter the amino acid sequence, whereas dS represents substitutions that do not affect the encoded amino acids. A lower dN/dS ratio indicates stronger evolutionary constraint and higher conservation, whereas a higher ratio suggests relaxed constraint or potential adaptive evolution. Statistical comparisons between NP and NPR groups were performed using the Wilcoxon rank-sum test (unpaired), and a P-value < 0.05 was considered statistically significant.

Additionally, the counts of human NP and NPR genes whose homologues have been identified in the model organisms in this study—including mouse, rat, fly, and nematode—were calculated and compared. This analysis shows which gene set—NPs or NPRs—better preserves its homologues across organisms. Data from these evolutionary conservation analyses are presented as bar graphs, with individual data points (dots) overlaid where applicable.

### Quantitative analysis of antisense RNAs, CpG Islands, and TATA boxes

To examine if there is a differential pattern in the distribution of genomic regulatory elements across NP and NPR genes, the presence of three major regulatory elements in their genomic loci was tested: antisense overlapping RNAs, CpG islands, and TATA boxes. For this purpose, the Ensembl gene IDs from all tested organisms—human, mouse, rat, fly, and nematode—were queried against the latest versions of their genomes in the UCSC Genome Browser (80). For the analysis and quantification of TATA boxes in NP and NPR genes, a genomic window of 50 nucleotides upstream of these genes was extracted from the UCSC Table Browser, using the human (hg38), mouse (mm39), rat (rn7), fly (dm6), and nematode (ce11) genomes as references. All unique NP and NPR gene IDs were inputted as a gene list, with parameters set to include only the window from -50 to +1 (i.e., 50 bp upstream of the transcription start site, TSS). These upstream sequences were downloaded and processed for TATA quantification using the ‘Elements Navigation Tool’ (ElemeNT – version 2023) (81). The number of identified TATA boxes for each gene was recorded, categorized into NP and NPR groups, and visualized using box and violin-dot plots. The presence and quantity of CpG islands and antisense transcripts were calculated by visual counting of these elements that overlapped the genomic locus of each gene.

### Epigenetic profiling of histone marks at NP and NPR loci

To test if NP and NPR genomic loci are subject to different epigenetic marks, quantitative analysis of major known regulators of histone modifications as indicators of poised promoter (H3K4me3), active enhancer (H3K27ac and H3K4me1), active transcriptional elongation (H3K36me3), and silenced chromatin (H3K27me3 and H3K9me3) was performed. For this purpose, three lineages of human cells including embryonic stem cells (ESCs), neuronal stem cells (NSCs), and differentiated neurons were examined for these modifications using their chromatin immunoprecipitation sequencing (ChIP-seq) data obtained from the ‘Encyclopedia of DNA Elements’ (ENCODE) database (82, 83). The human genome version GRCh38.p14 was used as the mapping reference, and genomic regions of -20 kb to +20 kb and -1500 bp to +200 bp relative to transcription start site (TSS) of each gene were considered as ‘enhancer’ and ‘promoter’ regions, respectively. Accession numbers of the samples used for this analysis are presented in the supplemental file S2A.

### Analysis of chromatin dynamics at NP and NPR loci

Four major chromatin states—bivalent, repressed, active, and inactive—were investigated in five different human tissues: colon, heart, liver, lung, and spleen. To analyze chromatin states, histone ChIP-seq datasets for each tissue were obtained from the ENCODE database in bigBed format. For each NP and NPR gene, the presence or absence of histone mark peaks within the gene-associated regions was determined based on overlap with the bigBed tracks. Genes were then categorized according to the detected combinations of histone marks, and the number of genes in each category was counted separately for each NP and NPR group. Their counts were converted into percentages and visualized as stacked bar plots. Statistical differences between NP and NPR groups were assessed using Fisher’s exact test. Samples for this analysis were obtained from ENCODE with their features and accession numbers listed in the supplemental file S2B.

### Transcription factor binding profiling

To identify the transcription factor (43) binding profiles of the *NPY* and *NPY1R* genes—representative of a ligand-receptor pair—shared data from two TF databases (ChIPBase and TFlink) were analyzed. ChIPBase utilizes data from ∼55,000 ChIP-seq experiments (84) and covers ∼3,000 regulatory factors (85). TFlink (https://tflink.net/) provides comprehensive TF interaction profiles for humans and six model organisms, including mouse (*Mus musculus*), rat (*Rattus norvegicus*), zebrafish (*Danio rerio*), fruit fly (*Drosophila melanogaster*), nematode (*Caenorhabditis elegans*), and yeast (*Saccharomyces cerevisiae*). TFlink data represent an integrated combination of both small-scale and large-scale experiments (86). To further evaluate the potential of *NPY* and *NPY1R* promoter regions to bind human TFs, TFBIND (https://tfbind.hgc.jp/) was employed, targeting the region 1,000 bp upstream and 1,000 bp downstream of the transcription start site (TSS) for each gene (Supplemental file S2C). TFBIND profiles are based on predictions using known human TF binding sites and are capable of assessing the binding potential of any DNA sequence (87). This combination of two experimental tools (ChIPBase and TFlink) and one prediction-based tool (TFBIND) was used for consistency and robustness. The identified TFs for *NPY* and *NPY1R* genes across ChiPBase, TFlink, and TFBIND, together with their shared (common) or unique subsets—derived from Venn analysis—are represented in the Supplemental file S3 (A–G).

### Gene set enrichment analysis (GSEA)

To gain insight into the biological processes and chromosomal regions linked to the TFs bound to the *NPY* and *NPY1R* genes, the ‘Database for Annotation, Visualization, and Integrated Discovery’ (24) was employed, using the ‘NPY TFs’ and ‘NPY1R TFs’ lists as input data (Supplemental file S3G), corresponding to the *NPY*- and *NPY1R*-specific TFs, respectively. Its output data are presented as tables, sorted by adjusted p-values, and include a full list of enriched genes for each hit.

### Statistics

Unless otherwise specified, statistical significance in this study was determined using two-tailed unpaired t-tests. Data are presented as Mean ± Standard Error of the Mean (SEM) in bar and violin plots, and p-values < 0.05 were considered statistically significant: *P < 0.05, **P < 0.01, ***P < 0.001, ***P < 0.0001, ****P < 0.00001, and *****P < 0.000001. Data were analyzed and presented via GraphPad Prism version 10.0.0 (GraphPad Software, Boston, MA, USA) for Mac. Figures were compiled and adjusted using Microsoft PowerPoint version 365 (Microsoft Corp.) and Adobe Illustrator version 2025 (Adobe Inc.).

## Data availability

All data are included within the manuscript, its figures, and the supplementary files.

## Author contributions

Conceptualization: WJK, HN, SSC, and SHJ. Methodology: HN, SHJ, SSC, and WJK. Validation: HN, SHJ, WJK, and SSC. Investigation: HN, SSC, SHJ, and WJK. Resources: SSC and WJK. Writing–original draft: HN, SHJ, and SSC. Writing–review & editing: HN, SHJ, WJK, and SSC. Visualization: HN, SSC, SHJ, and WJK. Supervision: WJK and SSC. Funding acquisition: SSC.

## Acknowledgments

The authors have no conflicts of interest to declare. We thank the genome sequence providers—including NCBI, UCSC, Ensembl, ENCODE, FlyBase, and WormBase—whose contributions to advancing comprehensive studies such as this are invaluable.

## Supplementary data captions

**Supplemental file S1:**

**A)** List of human neuropeptides (NPs, denoted by blue shades) and their cognate receptors (NPRs, denoted by red shades). Each NP may bind to several NPRs, and vice versa—one receptor may have affinities for different NPs. NP and NPR genes were identified via the International Union of Basic and Clinical Pharmacology (IUPHAR), and their names were then converted to official gene symbols and unique IDs using NCBI Entrez Gene. **B)** List of NP (blue columns) and NPR (red columns) orthologues with their species-specific gene IDs, used as the input datasets of this study. Human: *Homo sapiens*, Mouse: *Mus musculus*, Rat: *Rattus norvegicus*, Fly: *Drosophila melanogaster*, and Nematode: *Caenorhabditis elegans*.

**Supplemental file S2:**

**A)** Samples used in this study for analysis of epigenetic marks on NP and NPR genomic loci, obtained from ENCODE. **B)** Samples used for chromatin state analysis, utilizing human adult tissues examined with ChIP-sequencing (ChIP-seq) for various histone epigenetic marks. **C)** 1 kb upstream and downstream of the *NPY* and *NPY1R* genes (±1000 bp relative to the transcription start site, TSS), used for transcription factor binding site analysis in TFBIND.

**Supplemental file S3:**

**A)** Transcription factor (43) network of the *NPY* gene obtained from ChIPBase occupying the genomic window of 1 kb upstream and 1 kb downstream of the gene. **B)** Transcription factor (43) network of the *NPY1R* gene obtained from ChIPBase occupying the genomic window of 1 kb upstream and 1 kb downstream of the gene. **C)** Transcription factor network of the neuropeptide *NPY*, found by TFlink. **D)** Transcription factor network of the neuropeptide receptor, *NPY1R*, found by TFlink. **E)** TF network of *NPY* gene as predicted by TFBIND. Official gene symbols and descriptions were retrieved from the HUGO Gene Nomenclature Committee (HGNC), and gene list was finally formatted with one entry per row, represented as ‘final list’. **F)** TF network of *NPY1R* gene as predicted by TFBIND. Official gene symbols and descriptions were retrieved from the HUGO Gene Nomenclature Committee (HGNC), and gene list was finally formatted with one entry per row, represented as ‘final list’. **G)** TFs unique to the *NPY* gene (blue shaded), shared between *NPY* and *NPY1R* (dark grey shaded), or unique to the *NPY1R* gene (red shaded). Their regulatory effects on transcription levels are provided as quantitative results in the table.

**Supplemental file S4:**

**A)** The *NPY1R* gene exhibits a broader expression pattern compared to its ligand gene (*NPY*), which is linked to its unique genomic architecture and transcription factor (43) signature. **I-II)** While NPY protein significantly accumulates in the human central nervous system with lower expression in peripheral tissues, NPY1R protein is distributed across both neural and non-neural systems. This is denoted by red colors, with darker tones showing higher expression and lighter colors representing lower expression levels. Data was obtained from the Human Protein Atlas. **III)** A greater number of TFs regulate the *NPY1R* gene in comparison to its ligand gene, *NPY*. This result was confirmed using two experimental databases (ChIPBase and TFLink) and the TF prediction tool TFBIND. **IV–V)** The list of TFs linked to the *NPY* and *NPY1R* genes common to both ChIPBase and TFLink were analyzed using a Venn diagram and categorized into three classes: *NPY*-specific TFs, common TFs, and *NPY1R*-specific TFs. These three TF groups were then analyzed for their mode of action on transcription—activator, repressor, dual, or architectural. *NPY* TFs exhibit more binary activator and repressor—’ON/OFF’—modes, while *NPY1R*-specific TFs also include architectural TFs whose activity depends on gene and/or chromatin 3D structure.

**B)** There is a lack of significant expression correlation between *NPY* and *NPY1R* transcripts across different organs of the human body. Most tissues exhibit abundant expression of the *NPY1R* gene (the receptor), while the narrower expression of *NPY* (the ligand) results in a low correlation coefficient (r) value. Data were obtained from ChIPBase.

**Supplemental file S5:**

**A)** Gene set enrichment analysis for the transcription factors specific to *NPY* gene, via DAVID. **B)** Gene set enrichment analysis for the transcription factors specific to *NPY1R* gene, via DAVID.

